# Somatotopic Arrangement of Eight Distinct Skin Areas in the Human Primary Somatosensory Cortex Derived from Functional Magnetic Resonance Imaging

**DOI:** 10.1101/2020.06.22.164871

**Authors:** W. R. Willoughby, Kristina Thoenes, Mark Bolding

## Abstract

Blood oxygen level dependent (BOLD) functional magnetic resonance imaging (fMRI) was used to investigate cortical activity associated with peripheral tactile stimuli in a small cohort of healthy humans. MR-safe automated pneumatic stimulators modeled after the Wartenberg pinwheel were used to generate tactile stimuli at regular intervals on eight disparate areas of skin. The phase-encoded BOLD responses of voxels in the cerebral cortex were characterized by the maximal normalized cross-correlation coefficients at time delays between an idealized response and the measure time course. Overall at the group level, the somatotopic organization of the somatosensory cortex (SI) follows the accepted homunculus model, but a noticeable amount of variation was observed between individual study participants. The surface areas of cortical regions in SI activated by tactile stimulation of different body parts were calculated, giving an estimate of cortical magnification factors. Data collected with the participant actively attending the stimuli were compared to data collected before the attention task. No significant attention-related changes were observed in the somatotopic maps or in time courses of voxels well-correlated to stimuli.

## 1 Introduction

Sensory systems offer investigators good starting points for understanding the functional organization of the brain and large portions of the mammalian cerebral cortex are devoted to processing sensory inputs (Krubitzer and Seelke 2012). Once these systems are well understood, researchers have a solid foundation on which to base subsequent studies of systems related to more abstract cognitive functions, like higher reasoning, language, and memory. Initially, neuroscientists were forced to use highly invasive methods to probe the mechanisms of sensation, which limited the applicability of such studies to humans. [1–5] More recently, advances in functional neuroimaging over the past several decades have opened new avenues for investigating sensation and perception non-invasively [6–20]. Because of its non-invasive nature and because of the wealth of fundamental knowledge derived from homologous systems in other mammals and from humans undergoing surgical procedures, functional imaging of sensory systems currently provides the best foundation for understanding the functional organization of the brain in healthy humans.

A large body of work has been published that pertains to the evolution, structure, and function of the somatosensory nervous systems of mammals, including humans [1,3,5,17,21–28]. Much of what is known about functional localization of somatosensory cortex has been learned through lesion studies, electrical recording of neuronal activity through implanted electrodes, and direct electrical stimulation of exposed brain tissue of conscious subjects [1,5,21,22,24,29–31]. However, in the time since Wilder Penfield and his colleagues published the results of their intraoperative experiments in the 1930s, the cartoon of the cortical homunculus has become pervasive in the field of neuroscience, despite the seminal neuroscientists’ early warnings about reproducibility [1, 32]. Follow-up studies conducted since then have mostly supported the early findings of a somatotopic sequence in the Rolandic cortex [30], but there have been relatively few studies measuring individual, inter-subject differences in the functional anatomy of the somatosensory system [7, 9, 10]. Such data should offer fundamental insights into correlations between functional activity and each person’s unique macroscopic anatomy, as well as brain plasticity. Over the course of one’s life, changes in the makeup of the peripheral nervous system due to injury or other circumstances may be compensated for by corresponding changes in the central nervous system. The clinical implications of this avenue of research include streamlined preoperative neuroimaging and individualized treatments that would lead to higher success rates and improved quality of life for patients. Exploring individual variations in functional activation patterns due to somatic tactile stimulation is also a first step toward addressing deeper questions pertaining to neuroanatomical and cognitive evolution and development. We hypothesize that differences in the layout of functional localization in the somatosensory cortex between individuals can be probed non-invasively with functional MRI and an automated method for generating tactile stimuli on the surface of the skin.

Direct microelectrode recording and other invasive methods are not practical for routine study of the living human cortex. Human in vivo studies have been restricted by either a small pool of subjects eligible for intracranial operative measurements or by the technological limitations of less invasive neuroimaging techniques. EEG and MEG measure electromagnetic fields associated with neuronal activity and are known for their good temporal resolution, which can be less than 1 ms. However, these techniques are impaired by relatively poor spatial resolution on the order of 5 to 10 mm. Positron emission tomography (PET) and single-photon emission computed tomography (SPECT) have better spatial resolution, about 4 to 6 mm, but poor temporal resolution on the order of minutes along with the downside of requiring administration of small doses of ionizing radiation to subjects. Functional near-infrared spectroscopy (fNIRS) measures the attenuation of light with wavelengths between 700 and 900 nm and provides an estimate of the relative concentrations of oxy- and deoxy-hemoglobin in cortical tissue. The technique is limited by its poor spatial resolution and a limited penetration depth of about 3 cm. Functional magnetic resonance imaging (fMRI) is sensitive to changes in nuclear spin relaxation times due to paramagnetic effects of oxygenated hemoglobin on nearby protons. Like fNIRS, fMRI measures the effects of neurovascular coupling, which is indirectly related to neuronal activity. fMRI has good spatial resolution of about 1 to 2 mm, and advances in multiband pulse sequences have improved the temporal resolution of this technique to less than 2 s. The good spatial and temporal resolution makes fMRI our neuroimaging technique of choice for mapping individual differences in sensory somatotopic maps of human subjects.

To compare and contrast somatotopic maps generated with fMRI between individual humans and the accepted cortical homunculus, a pneumatic stimulation device was built that is safe to use in an MR environment. The device simulates the action of a Wartenberg pinwheel [33] over a small area of skin, but is made of plastic and does not puncture the skin or elicit pain. The device was fully automated and was synchronized to the acquisition of fMRI data, which enabled the use of analytical techniques based on stimulus timing and temporal correlation that have been used to study topographic maps of the visual and somatosensory systems [34, 35].

It is important to note that our method of mapping differs in fundamental ways from the methods originally used to reveal the cortical homunculus. Intraoperative experiments applied an electrical potential difference to somatosensory cortical structures with a resolution of about 2 mm. Qualitative data were then taken from awake patients undergoing surgery regarding their perception of sensation. In contrast, for our fMRI experiments, pressure changes at a certain frequency are applied to the surface of the skin, where they are transduced into electrochemical signals that are passed on to the somatosensory cortex. Then, quantitative data in the form of percent change in MR contrast as a function of space and time coordinates are recorded. The direction of input-output pathways are anatomically reversed, and the implications of these differences are not completely clear.

## 2 Methods

The protocol for this study was approved by UAB’s institutional ethics review board (IRB). Seven healthy participants (3 female, 4 male, ages 19-34) with no MRI contraindications or previous history of neurological disorders were recruited and gave their voluntary, informed consent to participate.

### 2.1 Pneumatic light touch stimulation

A pneumatic device was developed and built to automate and synchronize tactile stimulation. Light touch stimuli were produced by alternatingly pressurizing and depressurizing opposite ends of 8 plastic pneumatic cylinders (LEGO part ×189×01) and driving plastic pistons back and forth. Two small toothed wheels were attached to each of the 8 pistons. A photograph of one of the devices is shown in Figure 1. The stimulators were capable of delivering light touch to a 4 cm^2^ area of skin. Each cylinder was driven with a 5 port, 2-way valve connected using lengths of polyurethane tubing. The state of each valve was switched independently with a 24V solenoid using a microcontroller (Arduino MEGA) and 5V relays. Constant air pressure of 200 kPa was provided to each valve by an air compressor. The beginning of a stimulation sequence was triggered by a TTL pulse output from the MRI scanner at the start of fMRI volume acquisition, ensuring synchronicity between stimulus presentation and acquisition of functional data. Each stimulator was attached to the body using elastic bands, and self-adhesive bandages were used in more sensitive areas to prevent discomfort from pinching. Before each experiment and after being placed inside the bore of the scanner, the participant was asked to verbally announce which body part was being stimulated as a stimulation sequence was run. This was done to ensure that each stimulator was attached as intended and receiving adequate air pressure to induce sensation.

**Figure 1:**
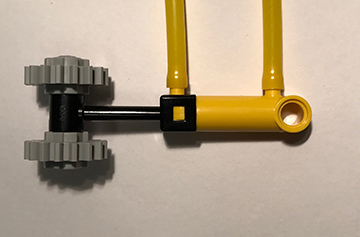
Photograph of one of the eight pneumatic tactile stimulators used for somatosensory topographic mapping experiments.

### 2.2 Paradigm

Figure 2 summarizes the traveling wave paradigm that was used to carry out the somatosensory mapping for each subject. Eight stimulators were attached to eight disparate body parts on the participant’s body, all but one of which were placed with left laterality, as shown in Figure 2 (a). Colored circles depict the locations of each of the stimulators: 1) the middle of the inferior side of the left foot, 2) the lateral side of the left shin, approximately halfway between the ankle and knee, 3) the lateral side of the left thigh, approximately halfway between the knee and hip, 4) the left lateral side of the torso at approximately the bottom of the rib cage, 5) the anterior side of the left forearm, approximately halfway between the elbow and wrist, 6) the distal phalanx of the left middle finger, 7) the left lateral side of the neck, at the approximate level of the C4 vertebra, and 8) the middle of the forehead, on the glabella.

**Figure 2:**
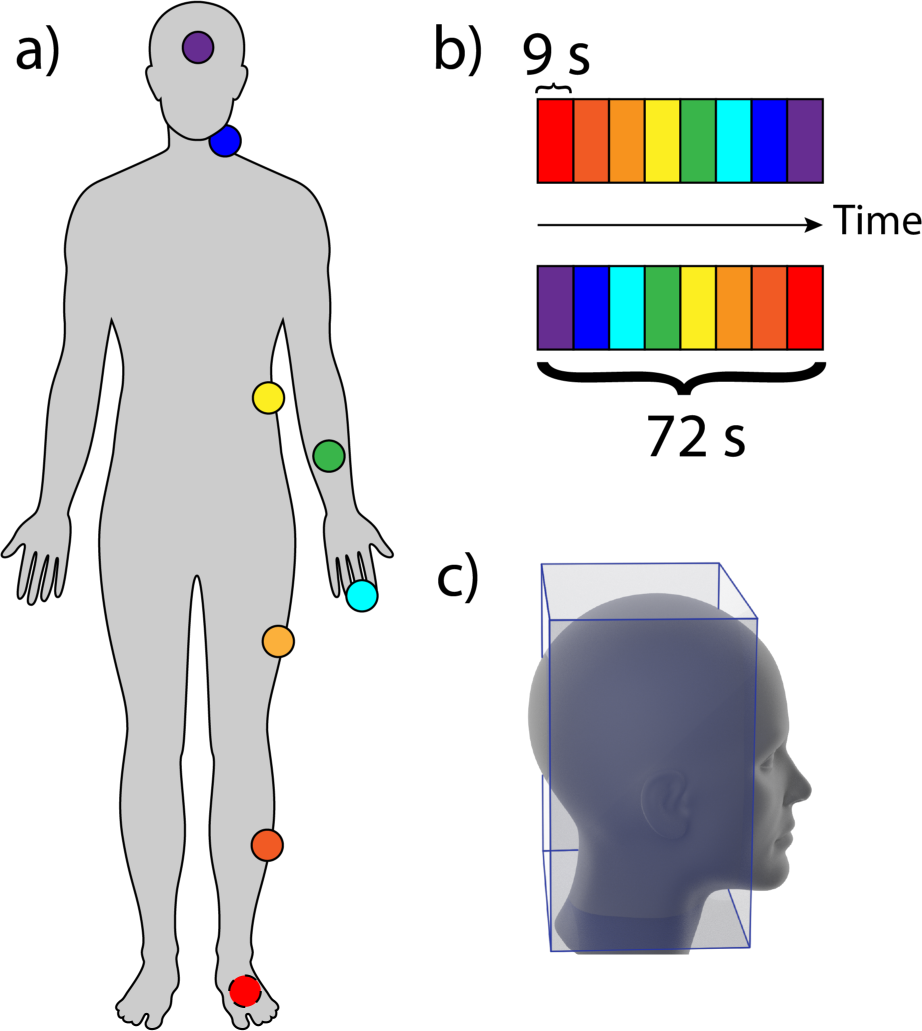
Stimulation paradigm for somatosensory mapping with fMRI. a) Colored circles represent approximate stimulator locations. b) Colored boxes depict timing of light touch stimuli. Top: forward sequence. Bottom: reverse sequence. Each stimulator was active for 9 s before switching to the next. Total stimulation period was 72 s. Each run consisted of 5 stimulation periods. c) Rendering of coronal functional imaging slab placement. Slices were stacked in the anterior-posterior direction. Phase encoding direction was right-left.

Figure 2 (b) illustrates stimulator timing for forward and reverse stimulation sequences. Each colored block represents the time interval over which the stimulator with matching color is active. Only one stimulator was active at any given time. The time axis runs horizontally from left to right. The top sequence was defined as the “forward” sequence, and the bottom sequence was defined as the “reverse” sequence. Data acquisition began after three TRs were discarded by the scanner to ensure the MR signal had reached steady state. The first stimulator began its 9 s cycle at the start of data collection at the beginning of the fourth TR. Each functional imaging run lasted for 5 stimulation cycles that were each 72 s, for a total of 6 minutes per scan. Four scans were acquired for each participant, two using the forward sequence (foot to head) and two using the reverse sequence (head to foot). For one forward and one reverse scan, the participant was asked to focus on the tactile sensation and press a button when the stimulus ended and switched to the next body part.

### 2.3 MRI acquisition

MR imaging was carried out at 3T using a Prisma scanner (Siemens Healthineers, Erlangen, Germany). A T1-weighted, magnetization-prepared rapid gradient echo (MPRAGE) 3D imaging sequence with 1 mm isotropic voxels was used to acquire high-resolution images for co-registration of EPI volumes and for cortical parcellation and surface reconstructions. Functional blood oxygen level dependent (BOLD) imaging was done using a T2*-sensitive, multi-band, single-shot 2D gradient echo echo-planar imaging (EPI) sequence (CMRR, University of Minnesota). The repetition time (TR) was 2 s, during which 50 contiguous 2 mm coronal slices were acquired. The echo time (TE) was 35 ms, and the flip angle was 75°. In-plane resolution was set by a 96 × 96 acquisition matrix and a 192 mm field of view (FOV), yielding 2 mm isotropic voxels. Phase encoding was in the right to left direction, and the multi-band acceleration factor was 2 with no in-plane acceleration. Five functional volumes with phase encoding in the right to left and left to right directions were acquired at the end of the scanning session for EPI distortion correction.

### 2.4 Data processing

EPI data was processed using AFNI (NIMH) and FreeSurfer (http://surfer.nmr.mgh.harvard.edu/) [36–38]. Cortical parcellation and generation of surface meshes with curvature maps were done using FreeSurfer. Slice timing alignment used AFNI’s 3dTshift function, and distortion correction was applied using the blip up-down technique built into the afni_proc.py script. BOLD time courses were detrended by subtracting third-order polynomials fit to the data voxel-wise by least squares regression. Geometrical alignment of fMRI volumes to high-resolution anatomical images was achieved using a signed local Pearson correlation cost functional. No spatial filtering was applied to the EPI data. Affine transformations between each subject’s anatomical scan and the MNI Colin 27 Average Brain template were calculated using linear interpolation between points. Each transformation consisted of 12 parameters and was used to map an individual’s functional data into a common template space. When no comparison between subjects was necessary, individual maps were produced in the subject’s native anatomical space. fMRI volumes were censored if motion was estimated to be greater than or equal to 0.3 mm. If more than 25% of volumes were censored, the entire dataset was discarded. Due to these exclusion criteria, one participant was not included in the analysis. Data from the remaining six participants were analyzed.

AFNI’s 3ddelay function was used to create voxel-wise estimates of temporal cross-correlation coefficients between the measured response of a voxel *r*(*t*) and an ideal response *s*(*t*) based on stimulus timing and a canonical hemodynamic response function (HRF). A gamma variate function

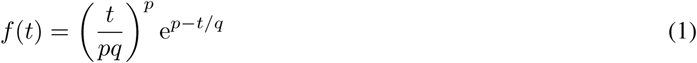

was used for the HRF, where *p* = 8.6, *q* = 0.547. *f* (*t*) was convolved with a boxcar function *b*(*t*) with duration 9 s, beginning at *t* = 0 s and repeating every stimulation period, *T* = 72 s, such that the model for an ideal response is

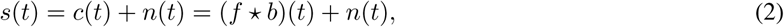

and the measured response is modeled as

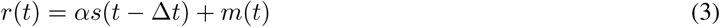

where *c*(*t*) = (*f* ⋆ *b*)(*t*) is the convolution of *f* (*t*) and *b*(*t*), *n*(*t*) and *m*(*t*) are noise components of the ideal and measured response function, respectively, is a scaling factor, and t is the time delay between the ideal and measured response. If it is assumed that the noise components are white noise with zero mean and are uncorrelated with each other and with *c*(*t*), then the cross-correlation *R*_*sr*_ between *s*(*t*) and *r*(*t*)can be expressed as a function of the time delay [38]

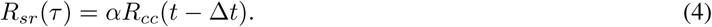

*R*_*cc*_(*t* − Δ*t*) is maximized when *τ* = Δ*t*, so calculating *R*_*sr*_(*τ*) and finding the maximum therefore gives the time delay Δ*t* and a normalized correlation coefficient *ρ* = *R*_*sr*_(Δ*t*)*/R*_*rr*_(0)*R*_*ss*_(0). To avoid reporting delays greater than 10% of each time series’ total length of 360 s, a second ideal time course was used to perform another cross-correlation analysis. This time course was delayed by *T/*2 = 36 s from the one starting at *t* = 0 s. This ensured that increases in delay variance due to the spectral characteristics of fMRI noise were mitigated [38]. The ideal time course beginning at *t* = 0 is plotted at the top of Figure [time courses].

Once delays and correlation coefficients were calculated for each functional scan, results from forward and reverse scans were combined to produce activation maps, one for which the participant was not attending the stimuli, and one for which the participant was asked to actively attend the stimuli. This was accomplished by combining the forward delays and correlation coefficients with respect to the ideal time course starting at *t* = 0 s and the reverse delays and correlation coefficients with respect to the ideal time course starting at *t* = 36 s. If a voxel was correlated to both forward and reverse delays, then the delay with the greatest correlation coefficient was used for that voxel. Voxels with correlation coefficients *ρ* ≥ 0.35 were divided into eight equally sized delay bins which corresponded to the body part being stimulated at that time.

Volumetric delay and correlation data were mapped onto each subject’s cortical surface using AFNI’s 3dVol2Surf function. Values from voxels lying on a line connecting adjacent nodes of the outer white matter surface and the pial surface were averaged to produce surface values. In order to accentuate cortical areas that were maximally correlated with stimulation of each body part, single cluster cross-correlation thresholds were found by incrementing the threshold until one voxel cluster in a region of interest (ROI) spanning the postcentral gyrus remained.

## 3 Results

Figure 3 shows the results of delay analyses for six participants when they were not attending the tactile stimuli. Each frame in Figure 3 is labelled with participant identifiers A-F. Inflated cortical surface maps show regions of negative curvature as dark gray areas and regions of positive curvature as light gray areas. Two views of the inflated surfaces are presented in the left portion of each frame, where the topmost perspective is a superior view, and the bottom perspective is a lateral view. The solid white lines on the surfaces depict the extents of the echo-planar imaging volume. Colors overlaid on the surfaces correspond to the body part with which BOLD time courses in that area were maximally correlated. The correlation threshold for the inflated surfaces was set at *ρ* ≥ 0.35.

**Figure 3:**
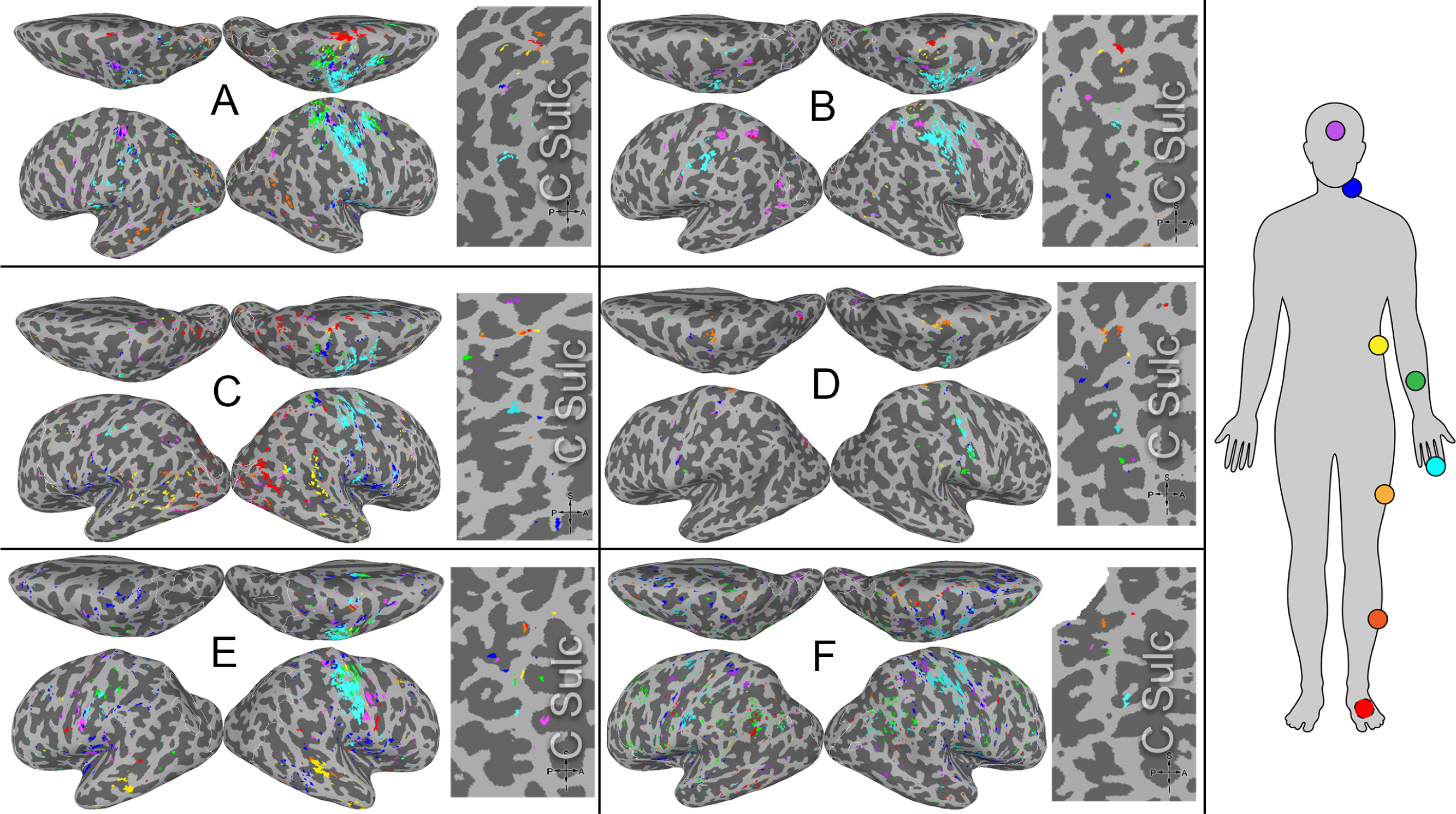
Single-subject results from delay analysis. A-F (left): Inflated cortical surfaces (areas of negative curvature, dark shaded regions; areas of positive curvature, light shaded regions) are overlaid with color-coded map of activation regions with cross-correlation coefficients *ρ* ≥ 0.35. A-F (right): Flattened cortical surface centered on ROI encompassing postcentral gyrus, extending from paracentral lobule to lateral sulcus and from central sulcus to postcentral sulcus. Correlation thresholds for each delay bin have been increased to yield one voxel cluster in the ROI that was correlated to each body part and emphasize areas of maximum correlation. Correlation thresholds are listed in Table [thresholds]. Right: Color key for activation correlated to stimulation of different body sites.

The right portion of each frame in Figure 3 shows a view of the flattened cortical sheet around an ROI that encompassed the postcentral gyrus of the cortical hemisphere that was contralateral to somatic stimulation. The ROI spanned from the paracentral lobule to the lateral sulcus, and from the anterior bank of the central sulcus to the posterior bank of the postcentral sulcus. This ROI was based on prior anatomical knowledge of the human primary somatosensory cortex [1, 30]. Correlation thresholds were independently adjusted for each body part in order to yield a single voxel cluster in the ROI and accentuate peak areas of activation. The thresholds are given in Table [thresholds].

Representative mean time courses from participant A are plotted in Figure 4. The top time series is the ideal time course initiated at *t* = 0 s. Each colored line beneath the black ideal series is an average of voxels within the ROI shown in the flattened maps of Figure 3. The color of each line corresponds to the delay bin those voxels were assigned to based on which time delay of the ideal series was maximally correlated to the BOLD response in that voxel. The delay analysis for the bottom four time courses in the plot, colored green, cyan, blue, and violet, corresponding to forearm, finger, neck and forehead stimulation, used an ideal time course that had been advanced by one-half of the stimulation period *T/*2 = 36 s. Each time course has the mean value subtracted and is vertically offset for clarity. No additional scaling was applied to the experimental data. The ideal time series was scaled to roughly the same scale as the experimental time courses. The dashed diagonal lines are all parallel and accentuate the time delay between peaks of the BOLD response in each time series. The colorbar at the top of the figure depicts the forward stimulation sequence used in the functional run during which these data were acquired.

**Figure 4:**
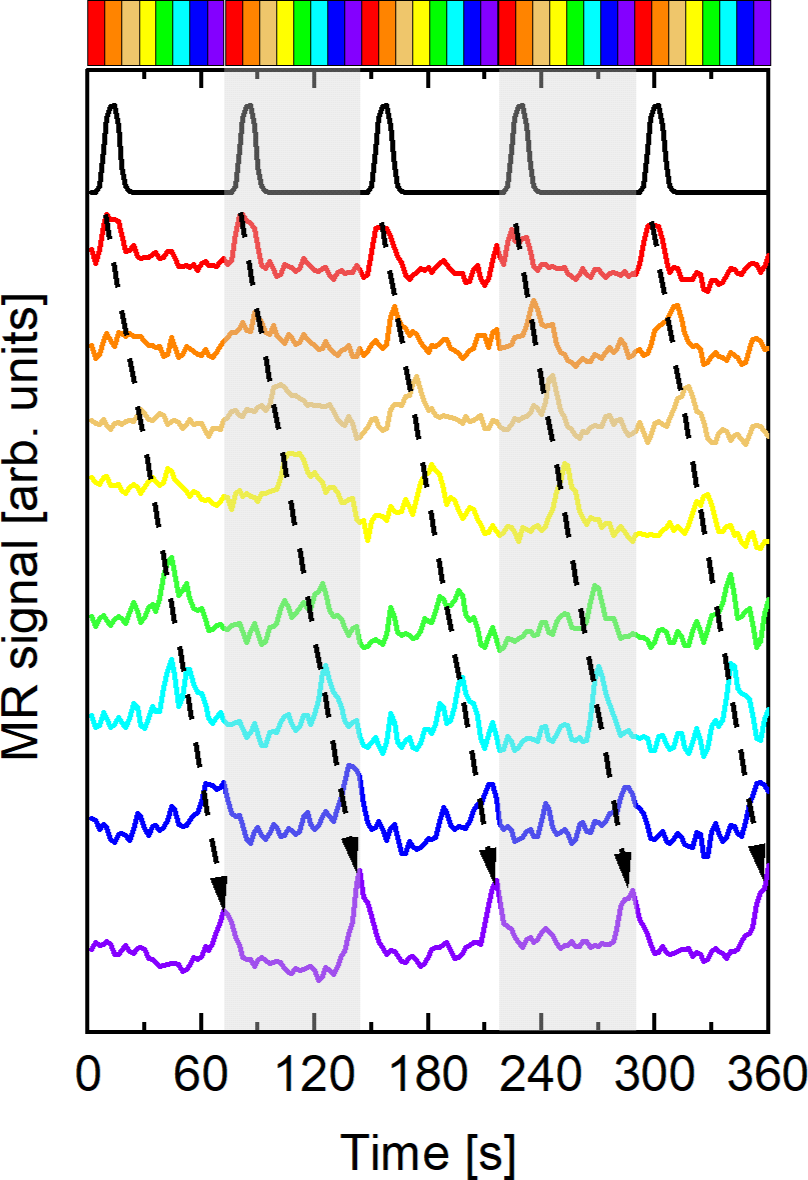
Representative mean time courses (subject A) averaged from voxels in ROI with *ρ* ≥ 0.35. Curves color-coded according to time delay, Δ*t*, with respect to the ideal time course (black curve). Time courses have been vertically offset for clarity. Top: Colorbar summarizing stimulus timing for the forward paradigm depicted in Figure 2. Dashed lines are parallel and are shown to provide a guide for the eye.

Overall, the maps show the somatotopic organization of S1 for each participant is roughly consistent with the accepted cortical sensory homunculus. Lower limb stimulation correlated strongly with the BOLD response from superior, medial areas of the postcentral gyrus and posterior paracentral gyrus contralateral to the stimulation. These areas are colored red and orange in the figure. Trunk stimulation near the left rib cage caused peak activation in different areas that was dependent on the individual, but peak activations, shown in yellow, were generally localized between the medial longitudinal fissure and the accepted hand area of S1 [39,40]. Stimulation of the left second finger was closely correlated to hemodynamic responses from this hand area. Peak responses, colored cyan in Figure 3, were at the anterior bank of the postcentral gyrus in participants A and F but were at the posterior bank in others. Stimulation of the anterior forearm produced responses near the accepted upper limb area in participants A, B, and E, but responses from participants C and F were localized in more superior and posterior regions of the postcentral sulcus and gyrus, respectively. Forearm stimulation was most correlated to BOLD responses in the most inferior regions of the postcentral sulcus, near the lateral sulcus, in participant D. Stimulation of the left side of the neck, close to the clavicle, and stimulation at the center of the forehead produced peak responses in the postcentral sulcus for each participant. Responses to head and neck stimulation in participants A and E were localized in the same area, about one-third of the way down the sulcus from the lower limb area. Forehead stimulation was in a similar area for participants B, C, and F, and neck stimulation was in a similar area for participant D.

From the peak correlation thresholds in Table [thresholds], it is clear that finger and forearm stimulation produced the most robust cortical responses that were the most closely correlated to the timing of these stimuli. However, there was a significant amount of intersubject variability in the peak correlation thresholds for different areas of stimulation.

The cortical magnification factor of each body area can be surmised from the maps with uniform thresholding overlaid onto the inflated cortical surfaces. Swaths of cyan representing fingertip stimulation are much larger than areas of other responses, which is consistent with previous studies that noted large degrees of somatosensory cortical magnification in the hands and fingers [refs]. Cortical magnification factors for each body part were quantified by calculating the total area of surface nodes that correlated with somatic stimulation at *ρ* ≥ 0.35 and are presented in Table [cortical mag]. Assuming that the area of skin stimulated at each site was approximately the same, comparing total surface area of activation is equivalent to comparing cortical magnification factors.

### 3.1 Attending task

Flattened cortical maps showing the same ROI as in Figure 3, which encompasses the postcentral gyri contralateral to stimulation, are shown in Figure 6. The top row shows maps acquired during the first two functional runs, during which the participant was not given instructions to attend the stimuli. The bottom row shows maps acquired during the second two functional runs, where each participant received instructions to attend the stimuli and give feedback via button responses when the stimulator switched sites.

**Figure 5:**
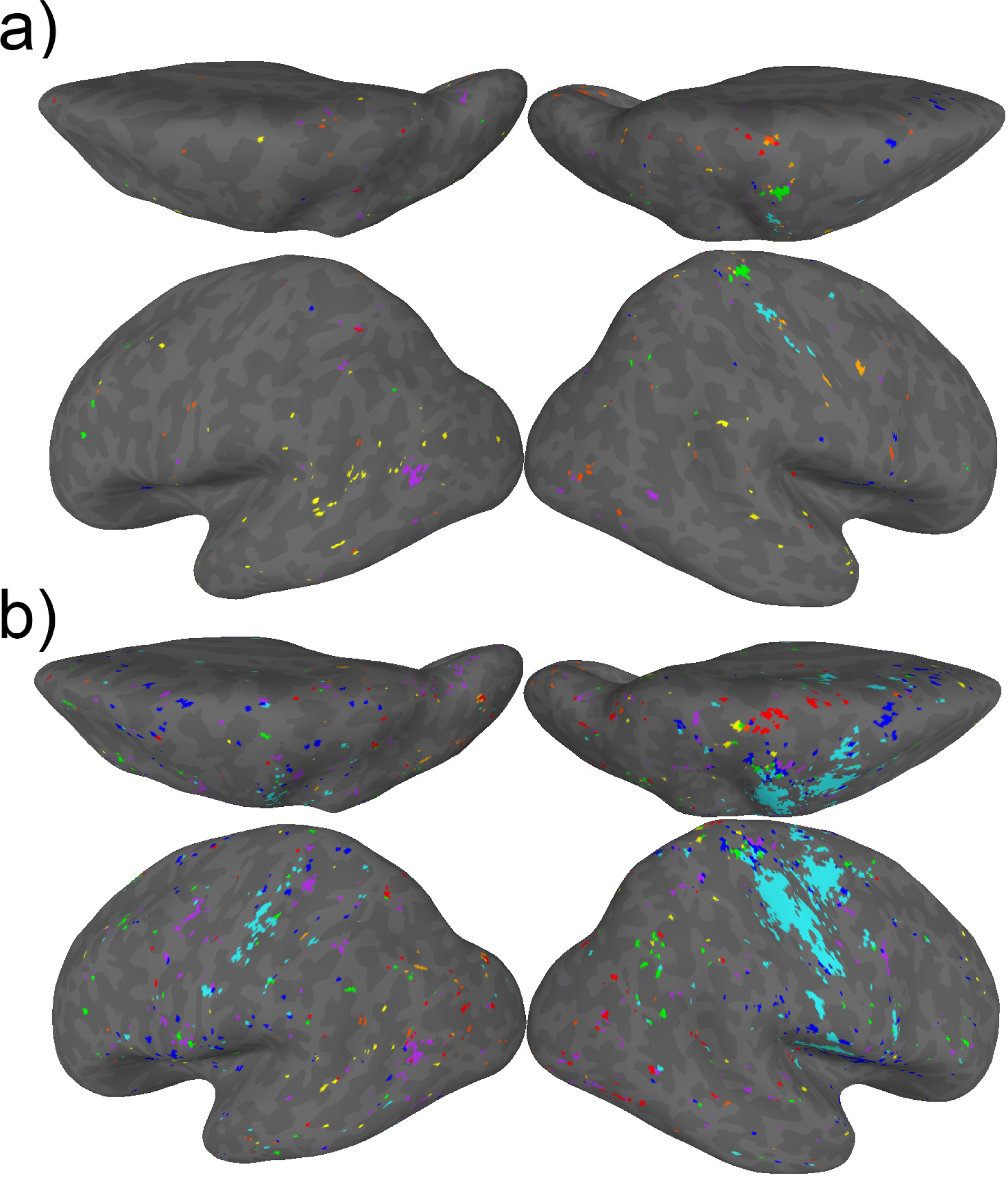
Concatenated group data obtained with no attention task on top of inflated cortical surfaces. (a) Obtained by averaging all individual subjects’ time series and performing delay analysis; correlation threshold set to *ρ* ≥ 0.35. (b) Obtained by superimposing individual delays and correlation coefficients using the thresholds from Table [thresholds].

**Figure 6:**
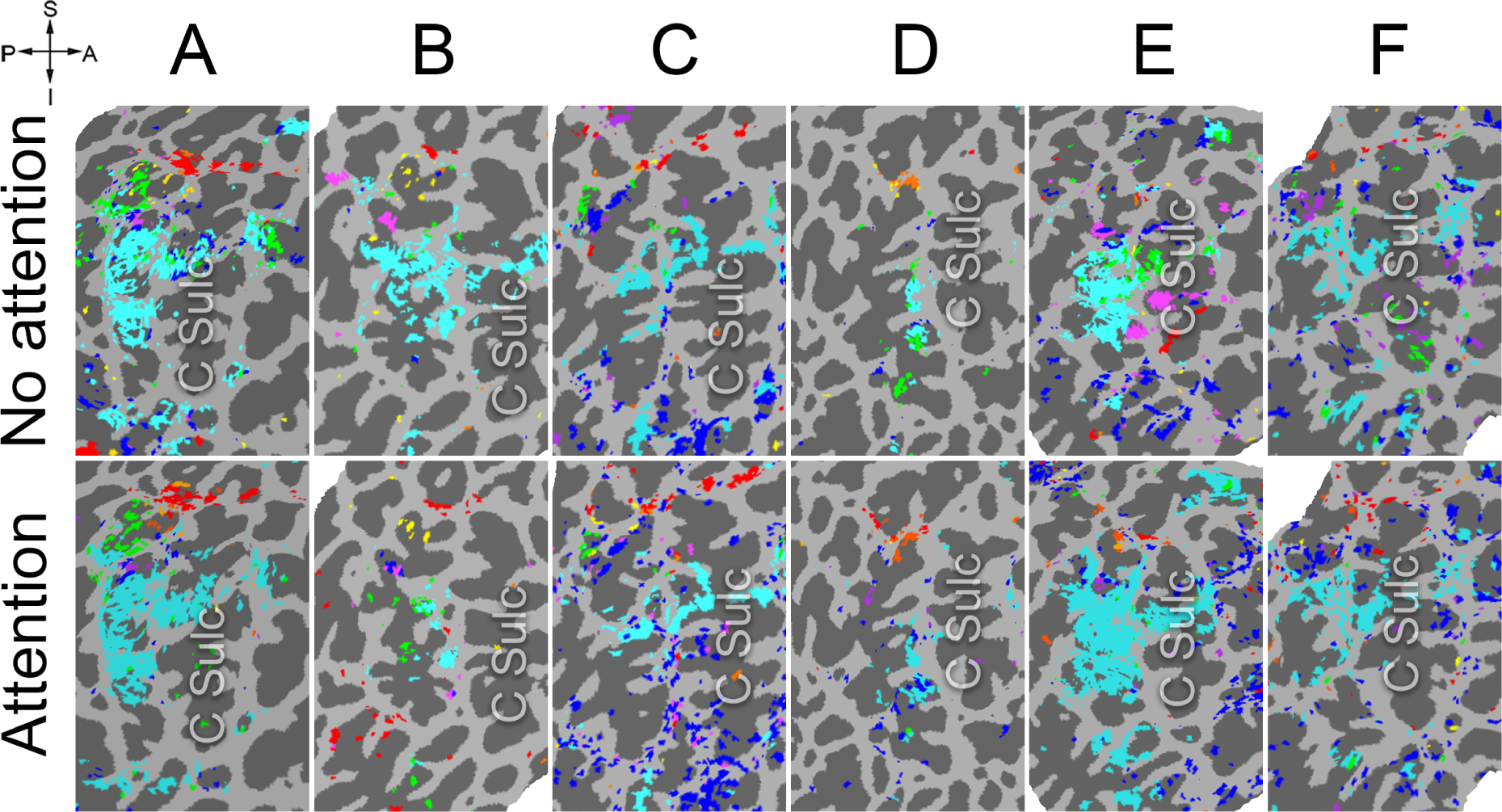
Flattened, individual cortical delay maps, threshold set to *ρ* ≥ 0.35. Top row: First two functional runs (forward and reverse stimulation sequence, no attention task). Bottom row: Second two functional runs (forward and reverse stimulation sequence, attention task). Top left: compass rose gives anatomical directions with respect to the maps. C Sulc: central sulcus.

Despite the noisiness of the maps at the threshold level of *ρ* ≥ 0.35, some variation can be seen between maps acquired while the participant was attending the stimuli. However, time courses from voxels showing significant activation did not vary markedly, both in terms of mean value and percent change, when the participant was and was not attending the stimuli.

## 4 Discussion

### 4.1 Large variability observed in individual somatotopic maps

For all of the subjects scanned, tactile stimulation of a body part correlated with the BOLD response in a specific region of the brain, which is consistent with the theory of functional localization as well as previous MRI-based somatosensory cortex mapping studies [7, 8, 11, 12, 26, 40]. The gross organization of the somatotopic map of each participant followed what has been known for decades: lower limb stimulation is associated with activity in the superior medial area of the postcentral gyrus, and peripheral stimulation of more superior body parts is associated with activity in the more lateral inferior areas of the cortex. However, the results presented in the previous section indicate that there is a large degree of variability in somatotopic maps across individuals, which is consistent with previous studies that noted individual differences in the layout of somatosensory cortex [7, 22]. Lower limb stimulation elicited BOLD responses that were the most consistent amongst the participants. Peak responses were near the midsagittal plane for most of the participants, often within 1 or 2 cm of the longitudinal fissure. Subject E demonstrated peak responses further down the gyrus, about 3 cm from the fissure. Trunk, distal arm, head, and neck stimulation generated localized responses that were more scattered amongst individuals. One major departure from the summary maps of Penfield and others is that, in several participants, stimulation of the head and neck areas correlated to activity in cortical structures more medial than those correlated to stimulation of the finger and torso.

The head area stimulator was attached midsagittal on the glabella. Voxels in the somatosensory cortex of the left and right hemispheres were correlated with stimulation in this area. For most of the participants, the forehead region of the somatosensory cortex lies deep within the postcentral sulcus in a region between the lower limb and upper limb area. This finding is consistent with a study on the facial representation on SI in macaques [21], as well as a recent fMRI study at 7 T which showed cortical activations in the same area upon cheek stimulation [35].

The group results shown in Figure 5 show many of the same features that are obvious in the individual maps. Stimulation of the fingertip activates a large swath of somatosensory cortex about halfway down the postcentral gyrus. Lower limb stimulation correlates strongly to medial areas of the cortex within 2-4 cm of the midsagittal plane.

The spatial distribution of peak cortical areas activated by stimulation of adjacent body parts, like the foot and leg, is also dependent on the individual participant. In subjects C and E, cortex correlated with foot stimulation is less than 1 cm from peak areas correlated with distal lower limb stimulation, while the other participants demonstrated a much larger spacing.

Another subject-dependent variable that was noticed was the cortical magnification factors of different areas of the skin. The distal middle phalanx elicited robust, widespread hemodynamic responses in the cortices of all of the participants. As seen in Table [cortical mag] the pial surface area of voxels that were correlated with0.35 to the finger stimulus timing was on the order of 10 cm2 for 4 out of 6 of the people scanned, nearly 25% of the total surface area of the postcentral gyrus and sulcus. For participant F, the surface area was about 15% less, while for participant D, the surface area was only 3 cm^2^. The cortical surface area correlated with the stimulation of other body parts also varied widely between individuals. It is worth noting that the cross-correlation thresholds used to obtain peak voxel clusters of activation and the total surface area of activated voxels seem to be correlated. This was expected and suggests that both may be considered, to some degree, a metric for cortical magnification.

There is some disagreement in the literature about how attention to tactile stimulation affects the cortical response [41–46]. Somatosensory cortical hemodynamic responses was not significantly different between trials in which the participant was or was not actively attending the stimuli. It is possible that any variation between maps is due to confounding factors between the two acquisitions.

### 4.2 Limitations

While care was taken to ensure each stimulator was attached at the same relative point on each participant’s body and was attached in the same way, it is possible that stimulator intensity for a particular body part differed between subjects leading to a source of intersubject variability in percent change of BOLD signal due to tactile stimulation. Also, during scanning, participants may have shifted and caused a change in relative stimulation intensity leading to a source of intrasubject variability. Additional experiments carried out in the same way on the same participant would have provided valuable information about this variability.

A deterministic sequence, either forward or reverse, was used for each of the functional imaging runs. No pseudorandom ordering was used, which may have affected the robustness of stimulation for later runs. Participant expectations and desensitization to stimuli may have had a confounding influence on the results, which may limit rigorous comparison of data collected in the same imaging session. Several studies, both fMRI and PET based, have investigated the effects of expectations on the processing of sensory input. One study by Drevets et al. suggested that anticipation of a localized tactile stimulus decreases cerebral blood flow in areas of cortex not responsible for processing input from the area in which stimulation is anticipated [47]. This implies that utilizing a predictable sequence for stimulation would result in decreased BOLD contrast due to reduction in blood flow throughout the somatosensory system.

### 4.3 Conclusion and Future Work

In conclusion, the results from these preliminary somatotopic mapping experiments are promising in that 3 T MRI was used to obtain maps with a resolution of about 2 mm. The automated stimuli allowed for the use of cross-correlation analysis to find areas of activation corresponding to the processing of somatosensory input from disparate body parts. Future studies using this technique will investigate whole body somatotopy at a finer scale, using more independent channels of stimulation, a more manageable and consistent method of attachment to the skin surface, and a pseudorandom sequence of stimulation that could increase BOLD contrast. Repeated trials on the same subject will also be carried out in order to estimate the intrasubject variability of the experiment.

## 5 Acknowledgements

This work was supported by funds from the UAB Department of Radiology Imaging Development Voucher Program.

---

**Table 1:**
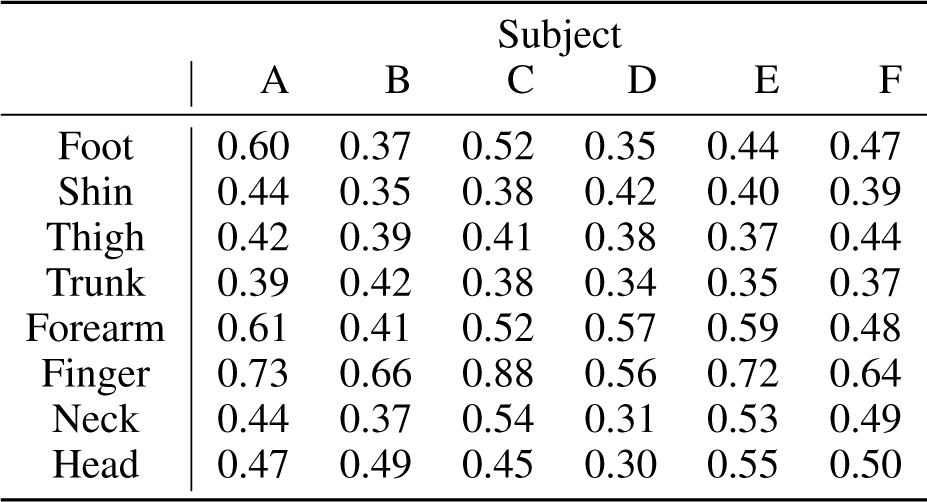
Single-cluster cross-correlation thresholds for flattened cortical maps shown in Figures 3 and 5 (b).

|  | Subject |  |  |  |  |  |
| --- | --- | --- | --- | --- | --- | --- |
|  | A | B | C | D | E | F |
| Foot | 0.60 | 0.37 | 0.52 | 0.35 | 0.44 | 0.47 |
| Shin | 0.44 | 0.35 | 0.38 | 0.42 | 0.40 | 0.39 |
| Thigh | 0.42 | 0.39 | 0.41 | 0.38 | 0.37 | 0.44 |
| Trunk | 0.39 | 0.42 | 0.38 | 0.34 | 0.35 | 0.37 |
| Forearm | 0.61 | 0.41 | 0.52 | 0.57 | 0.59 | 0.48 |
| Finger | 0.73 | 0.66 | 0.88 | 0.56 | 0.72 | 0.64 |
| Neck | 0.44 | 0.37 | 0.54 | 0.31 | 0.53 | 0.49 |
| Head | 0.47 | 0.49 | 0.45 | 0.30 | 0.55 | 0.50 |

**Table 2:** Cortical (pial) surface area [mm^2^] of voxels correlated to tactile stimulation with *ρ* ≥ 0.35.

|  | Subject |  |  |  |  |  |
| --- | --- | --- | --- | --- | --- | --- |
|  | A | B | C | D | E | F |
| Foot | 223 | 34.5 | 86 | 0 | 106 | 43.6 |
| Shin | 36.3 | 24.5 | 28.6 | 1.8 | 47.4 | 15.8 |
| Thigh | 44.6 | 40.9 | 17.4 | 91.7 | 4.4 | 5 |
| Trunk | 4.7 | 39.2 | 10.2 | 41.9 | 9 | 21 |
| Forearm | 126 | 65.8 | 74.2 | 250 | 361 | 187 |
| Finger | 1020 | 985 | 980 | 290 | 1070 | 840 |
| Neck | 213 | 6.2 | 346 | 66.7 | 308 | 263 |
| Head | 60.9 | 43.3 | 70 | 8.1 | 444 | 252 |

